# Asymptomatic infection and antibody prevalence to co-occurring avian influenza viruses vary substantially between sympatric seabird species following H5N1 outbreaks

**DOI:** 10.1101/2024.09.26.614314

**Authors:** Fiona Greco, Hannah M. Ravenswater, Francisco Ruiz-Raya, Chiara D’Avino, Mark A. Newell, Josie Hewitt, Erin Taylor, Ella Benninghaus, Francis Daunt, Gidona Goodman, David Steel, Jenny Park, Emma Philip, Saumya Thomas, Marek J. Slomka, Marco Falchieri, Scott M. Reid, Joe James, Ashley C. Banyard, Sarah J. Burthe, Emma J.A. Cunningham

**Author notes:** Shared last authorship (Sarah J. Burthe, Emma J.A. Cunningham).

## Abstract

Emerging infectious diseases are of major concern to animal and human health. Recent emergence of high pathogenicity avian influenza virus (HPAIV) (H5N1 clade 2.3.4.4b) led to substantial global mortality across a range of host species. Co-occurring species showed marked differences in mortality, generating an urgent need for better epidemiological understanding within affected populations. We therefore tested for antibodies, indicative of previous exposure and recovery, and for active viral infection in apparently healthy individuals (n=350) across five co-occurring seabird species on the Isle of May, Scotland, during 2023, following H5N1 HPAIV associated mortality in the preceding summer. Antibody prevalence to AIV subtypes varied substantially between species, ranging from 1.1% in European shags (*Gulosus aristotelis*) (to H5) to 78.7% in black-legged kittiwakes (*Rissa tridactyla*) (to H16 or both H13 and H16), and between 31-41% for three auk species (H5, H16 or both). At least 20.4% of auks had antibodies to an as yet unidentified subtype, suggesting further subtypes circulating in the population. We found low levels of active, but asymptomatic, AIV infection in individuals (1.6-4.5%), but excluded this as H5N1. Our results emphasise the importance of testing healthy individuals to understand the prevalence of co-circulating AIV subtypes in wild populations, and the potential for future reassortment events which could alter virus behaviour and impact.

## 1. Introduction

Emerging infectious diseases (EIDs) can substantially impact the health of humans, domestic animals, and wildlife, with severe economic and conservation costs^1^. Defined as pathogens that have recently appeared in a population or have rapidly increased in incidence or geographic range^2^, EIDs are largely driven by accelerated anthropogenic activity at the interface of wild and domestic species, with spillover of pathogens in both directions^3^. This is exemplified by the recent emergence of H5N1 clade 2.3.4.4b high-pathogenicity avian influenza virus (HPAIV), causing major mortality events in a wide range of both avian and mammalian novel hosts on a global scale^4,5^. Between 2021-2023, HPAIV caused the deaths of over 97 million birds worldwide, with over 250 wild avian species reported as newly affected with H5Nx HPAIV since 2021^6^. Severe impacts were noted in seabirds, previously unaffected by HPAIV^4,7,8^, and these events became a major conservation concern, with 30% of seabird species already listed as globally threatened^9^. Seabirds have been implicated in widespread viral transmission due to their global migratory routes increasing the propensity for spillover to a wider range of species^10,11^. With over 80 seabird species newly affected worldwide^6^, understanding how HPAIV may be circulating alongside low pathogenicity avian influenza viruses (LPAIVs) in these populations is critical to understanding the potential for viral recombination and spillover to other species.

AIVs naturally circulate in wild aquatic birds, distributed in *Anseriformes* (waterfowl) and *Charadriiformes* (including gulls, shorebirds, and terns) ^12,13^ worldwide. They are classified into different subtypes based on two glycoproteins occurring on the surface of the virus, namely Haemagglutinin (HA) of which there are 16 subtypes (H1-H16) and Neuraminidase (NA) of which there are 9 subtypes (N1-N9) (see Box 1 for key terms for avian influenza, and Box 2 for general terms). AIVs are defined as either LPAIVs or HPAIVs according to their infection outcome in domesticated *Galliformes* (poultry). Circulating AIVs within reservoir hosts are typically of low pathogenicity, however spillover of LPAIV H5 or H7 subtypes into high-density domestic poultry systems can facilitate the emergence of HPAIV following viral mutation within the HA glycoprotein^14,15^. While LPAIVs are typically associated with sub-clinical or mild infection in both wild and domesticated species^13^, the impact of HPAIV in wild birds can vary significantly. Some species succumb to debilitating infection whilst others tolerate and translocate infectious virus across vast distances^10,11,16^.

### Box 1

#### Glossary box of key avian influenza terms

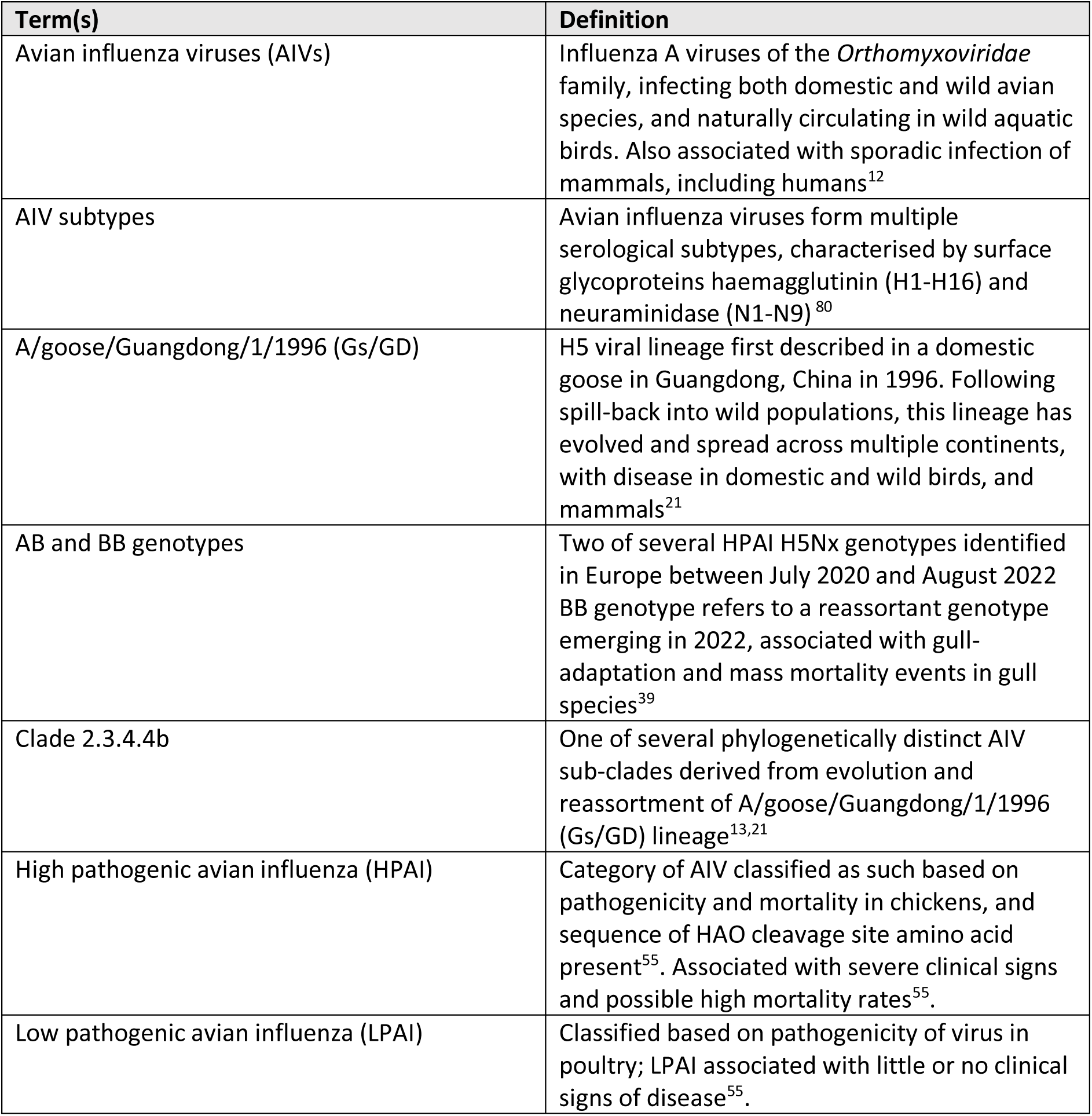

### Box 2

#### Glossary box of general terms

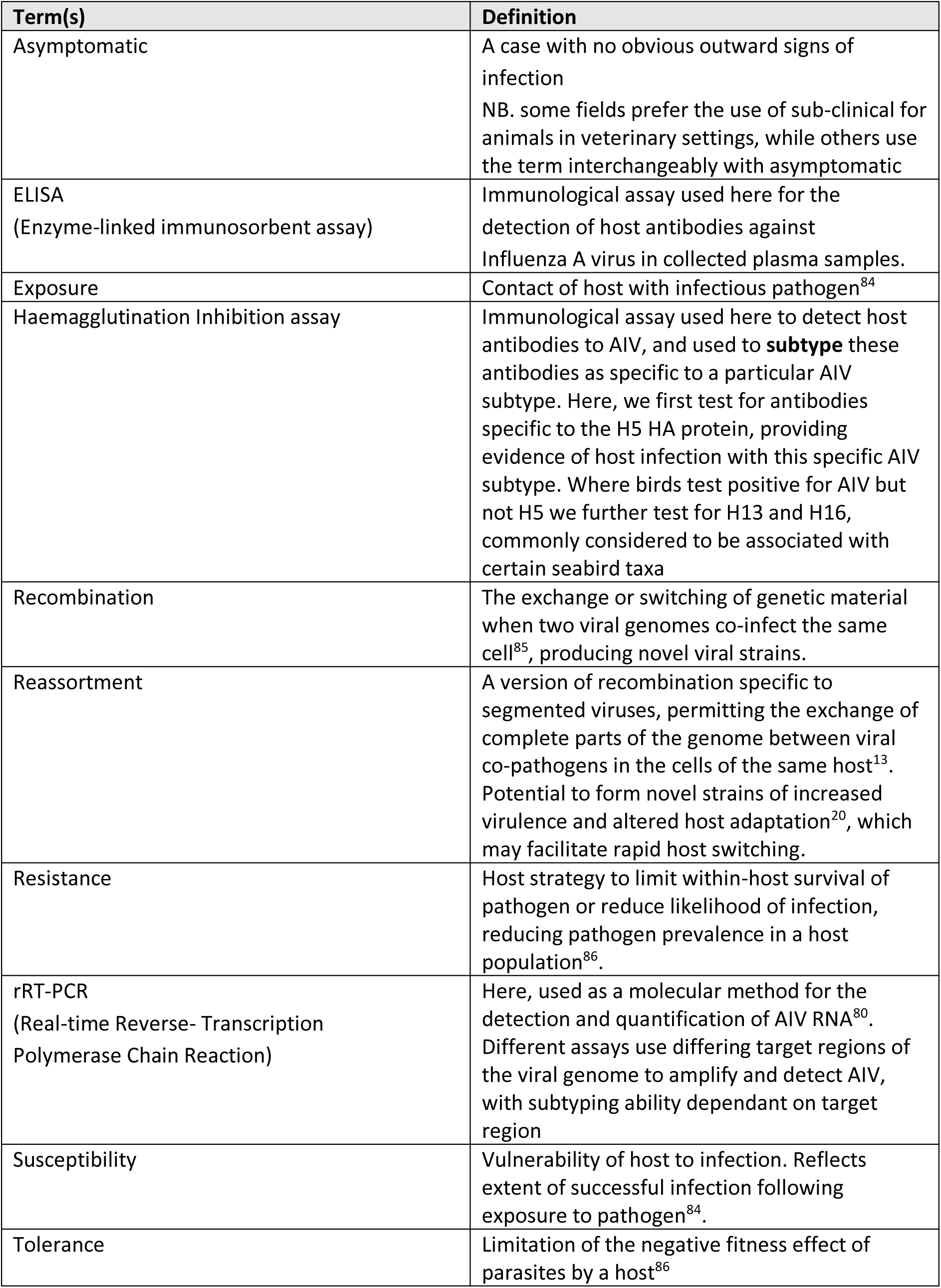

Along with genetic drift^17^, genetic reassortment of AIVs, potentially between HPAIV and regional LPAIVs^18,19^, can lead to the emergence of novel genotypes with the potential for altered virulence and host specificity^20,21^. The emergence of HPAIV subtype H5N1 in 1996, and subsequent evolution and reassortment of this A/goose/Guangdong/1/1996 (Gs/Gd) lineage has resulted in the presence of multiple phylogenetically distinct viral clades^22^, including H5N1 clade 2.3.4.4.b HPAIV^23^. Since 2020, this clade has become unique in its extensive geographical spread, disseminated by the seasonal migration of reservoir wildfowl species^24,25^, and associated with episodic outbreaks in poultry and wild birds^18,22,26^, with sporadic reports in humans^27^ and other mammals^28,29^. However, the recent scale and severity of H5N1 clade 2.3.4.4.b HPAIV has surpassed that of previous epizootic reports. Viral confirmation across 37 European countries^4^, and subsequent incursion into avian species in the Americas^30,31^, Africa^32,33^ and the previously unaffected sub-Antarctic and Antarctic^34^ highlights extensive, ongoing global impact. Reports of mortality in mammals caused by HPAIV, particularly in terrestrial carnivores and pinnipeds, and more recently including dairy cattle, span Europe and the Americas^4,28,29,35,36^, demonstrating an unusual impact upon non-avian species. Zoonotic potential is also of concern, with fifteen cases of H5N1 clade 2.3.4.4b HPAIV confirmed in humans between January 2022 and June 2024, varying from asymptomatic to fatal in outcome^37^.

The unprecedented nature of H5N1 clade 2.3.4.4b HPAIV necessitates an urgent global need to understand the ongoing epidemiology and implications of AIV in wild hosts. The life-history, foraging and migratory behaviour of seabirds makes them important potential disseminators of infection across broad spatial areas, particularly in long-distance migrants traversing global hemispheres^10,11^. Extreme diversity in species characteristics across seabirds also provides a challenge in predicting the dynamics of HPAI within this previously little affected species group. In addition, species variation in exposure to other AIV subtypes, including LPAIVs, could influence the susceptibility to, and outcome of HPAIV between host populations, and may determine species-specific routes for viral reassortment and the emergence of novel strains with altered biological characteristics. As an example, the Laridae family (including gulls and terns) are particularly noted for the circulation of H13 and H16 LPAIVs^38^ and recent evidence indicates viral reassortment between a gull-associated H13 LPAIV and H5N1 to produce a novel H5N1 genotype^39^. This genotype is believed to be particularly adapted to gulls and related taxa^39^, with extensive circulation noted in European herring gulls (*Larus argentatus*) ^4,39^. Knowledge of LPAIVs circulating in other seabird hosts remains comparatively limited.

While field evidence indicates marked species variation in mortality^7^, we know little of species differences in exposure, susceptibility or resistance to HPAIV, the potential LPAIV subtypes they may also be exposed to, or of their role in the spatial dissemination of infection^40^. Much of the existing data on HPAI comes from the testing of carcasses, which has limited scope for understanding variation in infection and immunity, for assessing capacity for survival and recovery, or for determining maintenance of virus in apparently healthy populations. It is therefore crucial to source epidemiological evidence for the prediction of ongoing viral transmission, persistence, and consequences of H5N1 clade 2.3.4.4b HPAIV among live populations. To this end, it is important to identify whether apparently healthy individuals have remained unexposed to infection, act as asymptomatic hosts, or have acquired some degree of immunity to reinfection following exposure and recovery. In addition, providing evidence of the range of AIVs concurrently circulating in these same populations beyond H5N1 clade 2.3.4.4b HPAIV is critical in predicting potential reassortment between different AIVs^19,39^, and in considering the possibility of at least some protective immunity acquired following prior infection with circulating LPAIVs^41,42^.

Here, we provide critical evidence of infection, exposure and recovery to AIV, and H5N1 clade 2.3.4.4b HPAIV in particular, within a multi-species assemblage of breeding UK seabirds. To do so, we examined five sympatric seabird species at an internationally important seabird breeding colony in the North Sea, where H5N1 clade 2.3.4.4b HPAIV was first confirmed during a high mortality outbreak event during the 2022 breeding season^43^. We first tested for the possibility of asymptomatic infection by testing for viral prevalence across all five species. We then tested for levels of exposure and recovery to HPAIV H5N1 and how this varied between species by testing for antibodies indicative of previous infection. Finally, we tested for antibodies to other LPAIVs known to occur in seabirds that may be co-circulating with H5N1, to establish the potential for reassortment which could further alter virus behaviour and impact.

## 2. Material and Methods

### 2.1 Study site

This study was conducted on the Isle of May National Nature Reserve (IOM), Scotland (56°11’N, 02°33’W), during the seabird breeding season (May-July) of 2023. Designated as a Special Protection Area (SPA), the IOM and surrounding islands comprise an internationally important assemblage of both cliff- and ground-nesting seabird species^44^. The IOM is ∼1.5km in length and 0.5km wide, and supports multiple species. Of those breeding in significant numbers, five species are listed on the UK Birds of Conservation Concern Red List^45,46^, while six others are considered of amber concern (Supplementary Table S1).

Bird mortalities from HPAIV H5N1 clade 2.3.4.4b were first confirmed on the IOM during the breeding season of 2022. Temporary suspension of research by statutory bodies necessitated a cessation of all seabird handling. Monitoring was restricted to distant observations and the limited testing of carcasses, with AIV detected via post-mortem swabs in several species: Atlantic puffin (*Fratercula arctica*), common eider (*Somateria mollissima*), black-legged kittiwake (*Rissa tridactyla*) and Arctic tern (*Sterna paradisaea*) ^43,47^. Symptomatic individuals were noted within the kittiwake, eider and herring gull populations, and additional mortality from HPAI suspected in these species. Mortality was particularly elevated in kittiwakes, with abnormally high mortality rates reported across the season (n=312 observed on land) (*NatureScot, unpublished data*). Clinically affected individuals and mortality from HPAIV H5N1 were also noted at other seabird colonies within the Firth of Forth and adjacent Lothian coast, highlighting the circulation of HPAIV within the region in 2022. The presence of HPAIV H5N1 was again confirmed on the IOM during the summer of 2023. This occurred after our sampling period (Supplementary material - Mortality Events).

### 2.2 Sample collection

All samples were collected by appropriately trained individuals, acting under UK Home Office Project License PEAE7342F in accordance with the Animals (Scientific Procedures) Act 1986, and under NatureScot Research License, while adhering to strict biosecurity and personal protective protocols.

To provide baseline epidemiological evidence of current and past AIV infections, samples were collected from apparently healthy individuals of five species from May to July 2023 (Table 1.). Species comprised European shag (*Gulosus aristotelis*) (n=100), black-legged kittiwake (n=100), puffin (n=50), common guillemot (*Uria aalge*) (n=50) and razorbill (*Alca torda*) (n=50). All species comprised breeding adults, captured at their breeding site, except for puffins which were caught in flight and where the reproduction status of 44% of sampled individuals was unknown. Capture occurred during late incubation or chick-rearing, and occurred across multiple sub-colony sites to reduce geographical bias in virus exposure and repeated site disturbance (Figure 1b.) All visible colonies were monitored post-sampling and where nests could be observed directly, all sampled birds returned to normal breeding activity following release.

**Figure 1.**
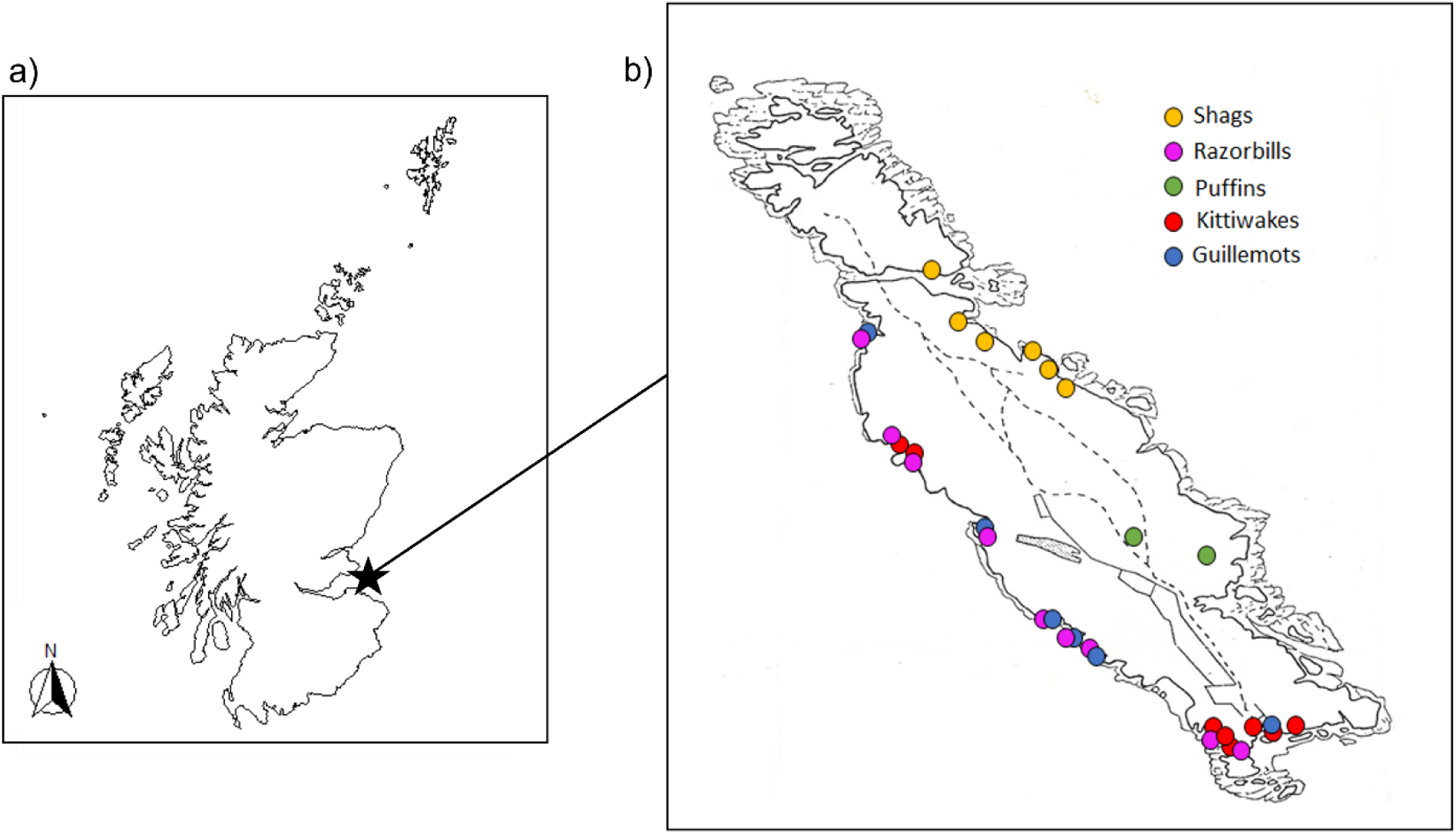
**a)** Location of study site - Isle of May NNR, south-east Scotland, UK (56°11’N, 02°33’W) **b)** Sub-colony sample sites for each species on the IOM. Sampling was spread across accessible sub-colony locations. Total sampling locations were as follows: puffins n=2, shags n=6, razorbills n=9, guillemots n=6, kittiwakes n=8. Not all available species and colony sites could be sampled and broader species assemblages occur at all sites across the island. ^48,49^

**Table 1.** Timing of sampling for each species.

| Species | Sampling dates |
| --- | --- |
| European shag (*Gulosus aristotelis*) | 29<sup>th</sup> May – 12<sup>th</sup> June 2023 |
| Black-legged kittiwake (*Rissa tridactyla*) | 25<sup>th</sup> May - 14<sup>th</sup> June 2023 |
| Puffin (*Fratercula arctica*) | 11<sup>th</sup> June – 6<sup>th</sup> July 2023 |
| Common guillemot (*Uria aalge*) | 21<sup>st</sup> June – 22<sup>nd</sup> June 2023 |
| Razorbill (*Alca torda*) | 10<sup>th</sup> June – 22<sup>nd</sup> June 2023 |

Unless already present, all individuals within the study received a unique metal BTO ring. To facilitate antibody testing for prior exposure to AIV, a volume of 0.5-1ml of blood was collected from either the brachial (shags, kittiwakes, guillemots, and razorbills) or medial tarsal vein (puffins) and placed in a pre-heparinised sample tube (∼8µl of heparin). These sampling volumes related to a maximum of 1.2-6% circulating blood volume obtained on a single occasion, based on reports of average species weight. To test for AIV shedding from either the oropharynx or intestinal tract, two swab samples were taken from both the cloacal and oropharyngeal regions using sterile, polystyrene viscose-tipped swabs. Swab ends were placed in sterile collection tubes. All tubes were uniquely identified and stored in a cool bag containing ice packs while in field prior to processing.

Blood samples were centrifuged at 3000rpm for 10 minutes no more than 120 minutes post-sampling, and separated into red blood cells and 80 µl plasma aliquots. Samples were stored at 4°C and either sent to the Animal and Plant Health Agency (APHA) for diagnostic assessment for avian influenza virus within seven days of sampling, or frozen immediately post processing at −80°C.

### 2.3 Diagnostic evaluation

Samples from a total of 350 individuals were collected for AIV diagnostics, however, not all assays could be performed for all individuals due to sample volume limitations or other sample requirements.

#### 2.3.1 RNA Extraction and rRT-PCR: detection of active viral infection

To detect and subtype active AIV infection, oropharyngeal (OP) (n=307) and cloacal (C) (n=309) swabs were first processed in Leibovitz’s L-15 Medium (ThermoFisher Scientific). Nucleic acid was extracted from samples using the MagMAX CORE Nucleic Acid Purification Kit (ThermoFisher Scientific) following the robotic Mechanical Lysis Module (KingFisher Flex system; Life Technologies) as described previously^50^.

Samples were tested for viral RNA via real-time reverse-transcription-PCR (rRT-PCR) ^50^. A minimum of two rRT-PCR assays were employed for each sample; an M gene screening rRT-PCR, targeting the conserved matrix protein (MP) segment of the viral genome, to detect influenza A virus RNA of all sixteen AIV HA subtypes^51^, and a high pathogenicity H5 specific rRT-PCR (H5HP rRT-PCR) used to define HPAIV as clade 2.3.4.4b H5Nx^52^, via a target region specific for clade 2.3.4.4b H5Nx viruses.

For both assays, a positive threshold cut-off of quantification cycle value (Cq) 36.0 was implemented^53^. Cq values below or equal to this value were classified as positive for AIV RNA while Cq values >36.0 were considered borderline and warranted further investigation. Failure to detect viral RNA through all 40 thermal cycles indicated a negative result.

Where possible, a third assay targeting the N1 gene (N1 rRT-PCR) was employed^54^ when borderline Cq results were reported for one, or both, of the previous rRT-PCR assays or where only one of the previously employed assays was positive (Supplementary Table S2) (n=11).

#### 2.3.2 Competitive ELISA: generic testing for antibodies to AIV nucleoprotein

Plasma samples from 285 individuals, across five species, were tested for avian influenza antibodies via a commercially available competitive ELISA (cELISA) kit (ID Screen® Influenza A Antibody Competition Multi-species, ID Vet, Montpellier, France). This kit detects antibodies against the nucleoprotein (NP) of Influenza A virus, highly conserved among all AIV subtypes, thereby detecting antibodies to all AIV subtypes H1-H16. Following manufacturers guidelines, a competition percentage (S/N%) of less than 45% was interpreted as positive for AIV antibodies. This value was calculated as follows:

S/N% = (OD_sample_/OD_NC_) x 100 where OD_sample_ is the optical density reading of the sample, and OD_NC_ is the mean optical density value of the negative control.

Assays were conducted at either APHA or the Ashworth Laboratories, University of Edinburgh. Thirty three percent of cELISA assays (95/287) were conducted at both sites, allowing verification of assay results across laboratories.

#### 2.3.4 Haemagglutination Inhibition assay: specific testing for antibodies to H5 subtype

Plasma samples from 289 individuals were tested via haemagglutination inhibition (HI) assay for the detection of H5 subtype-specific influenza A antibodies. An HI test antigen homologous to that responsible for the current outbreak was used (H5N1 HP (A/chicken/Wales/053969/2021)).

Inhibition of haemagglutination at a reciprocal HI titre of 16 or greater (i.e., serial dilution 1:16) was considered positive for H5-specific antibody^55^. Haemagglutination inhibition at subthreshold serial dilution titres of 1:8, 1:4 or 1:2 was considered a negative/weak reactor result.

#### 2.3.5 Haemagglutination Inhibition assay: specific testing for antibodies to H13 and H16 subtypes

To assess for previous exposure to H13 and H16 LPAIVS, both commonly circulating in members of the Laridae family^56^, we further tested a subset of individuals testing positive for antibodies via cELISA for antibodies specific to H13 and H16 LPAIV subtypes where plasma volumes permitted. This resulted in the testing of 21 kittiwakes, 13 puffins, 15 guillemots and 13 razorbills. HI assays were performed using test antigens H13N6 (A/gull/Maryland/704/1977) and H16N3 (A/gull/Denmark/68110/2002), with a reciprocal titre of 16 or greater indicative of a positive result.

#### 2.3.6 Competitive ELISA: specific testing for antibodies to H5 subtype

Eighty-four individuals testing positive for generic AIV NP antibodies were tested via competitive ELISA for the specific detection of antibodies against the haemagglutinin H5 of the Influenza A virus using the newly developed ID Screen® Influenza H5 Antibody Competition 3.0 Multi-species ELISA, ID Vet, Montpellier, France – following ‘Duck’ protocol. Following manufacturers guidelines, a competition percentage (S/N%) of less than 50% was interpreted as positive for antibodies against H5 haemagglutinin, with a S/N% between 50-60% classified as ‘doubtful’. Samples from seventeen individuals negative via cELISA for antibodies to all subtypes H1-H16 were included as negative controls.

## 3. Results

### 3.1 rRT-PCR - detection of active viral infection

Five out of 309 individuals tested positive for the presence of current AIV infection; four were detected via M-gene assay (3/44 guillemots (6.8%) and 1/45 puffins (2.2%)), and one via the H5 HP assay (1/48 razorbills (2.1%)). One guillemot that tested positive via the M-gene assay also tested positive for N1 specific RNA but no individual tested positive concurrently via the H5 and N1 assays. Both positive and borderline results are summarised in the supplementary material and combined gave lower and upper estimates of 1.6-4.5% AIV positive birds across the entire set sampled (Supplementary tables S2 and S3). Of the five individuals testing positive for AIV nucleic acid, three were detected on oropharyngeal swabs, and two on cloacal swabs. No individual tested concurrently positive on both cloacal and oropharyngeal swabs.

### 3.2 Serology – generic testing for antibodies to AIV nucleoprotein

Generic AIV antibody testing revealed previous exposure to AIV in all species tested, however, the percentage of positive individuals varied substantially across these species (range 1.1-78.7%) (Table 2, Figure 2). Shags had the lowest AIV antibody prevalence (1.1% of birds tested) and kittiwakes the highest (78.7% of birds tested) with all three auk species intermediate between these values (guillemots 41.3%, razorbills 31.9% and puffins 36.3%).

**Figure 2:**
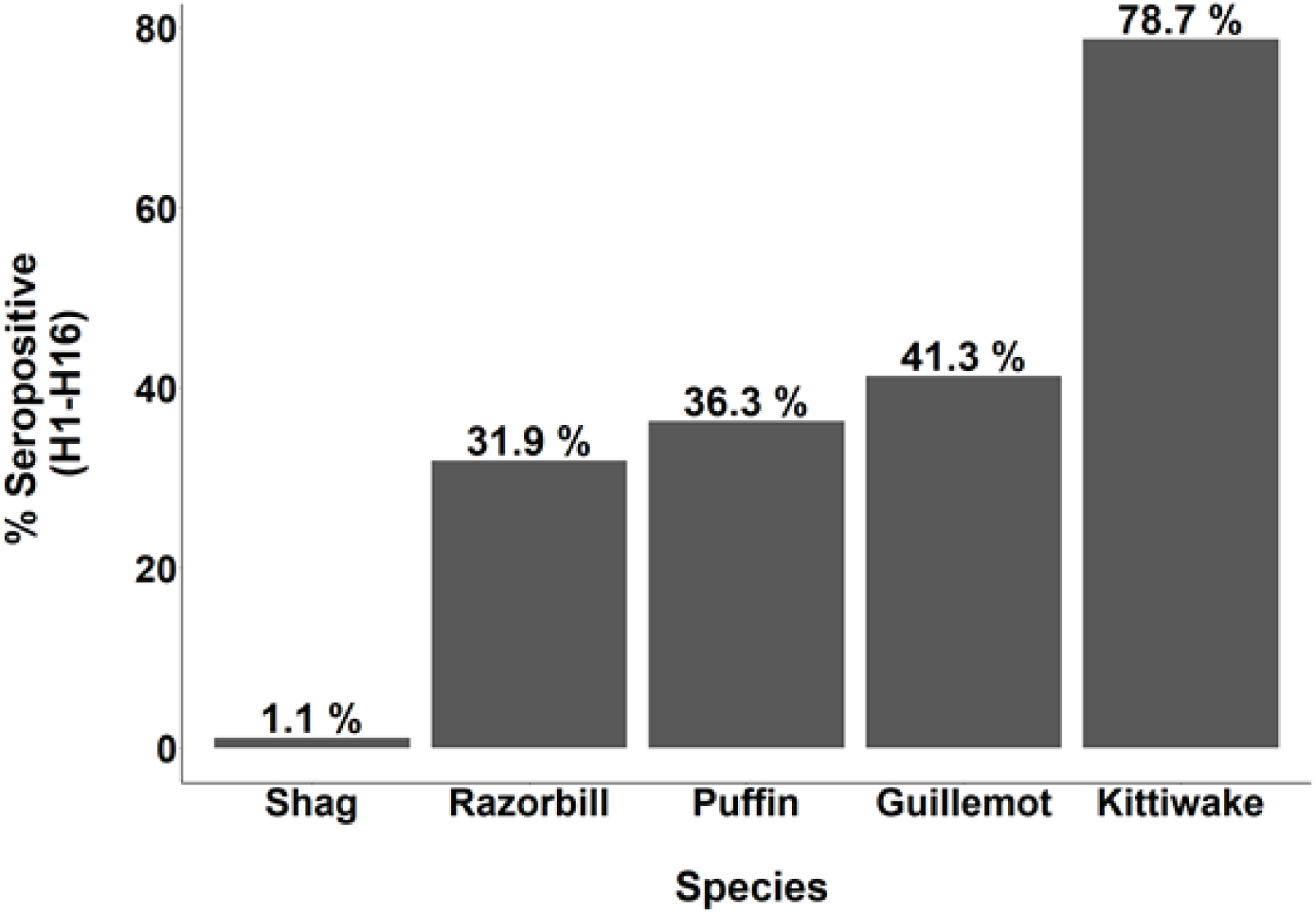
Between-species variation in seropositivity for generic antibodies to all AIV subtypes H1-H16 using ID Screen® Influenza A Antibody Competition Multi-species ELISA, ID Vet, Montpellier, France. Seropositive results range from 1.1-78.7% across species, with seropositive value for each species provided above each species bar.

**Table 2.** Percentage of positive antibody detections via competitive ELISA (for the generic detection of antibodies to AIV nucleoprotein) and via haemagglutination inhibition assay (for the detection of H5, H13 and H16 subtype-specific AIV antibodies) for individuals of each species. Number of positives in those tested is given in brackets. Incidences of co-occurring antibodies to more than one subtype, and number of individuals in which this occurred also given. *H13/H16 HI assays were only performed in individuals testing positive via cELISA, and negative via H5 HI, with sufficient plasma remaining. Note, the single shag positive by cELISA was volume depleted and could not be assessed for H13/H16.

| Species | cELISA<br>(H1-H16) | H5 HI | H13 HI | H16 HI | H5 ELISA | Antibody co-occurrence |
| --- | --- | --- | --- | --- | --- | --- |
| Shag | 1.1%<br>(1/87) | 0%<br>(0/92) | Not<br>tested* | Not<br>tested* | **100%**<br>**(1/1)** |  |
| Kittiwake | 78.7%<br>(48/61) | 0%<br>(0/69) | **19.0%**<br>**(4/21)** | **95.2%**<br>**(20/21)** | 0%<br>(0/39) | **H16 + H13 (n=4)** |
| Razorbill | 31.9%<br>(15/47) | 0%<br>(0/44) | 0%<br>(0/13) | **30.8%**<br>**(4/13)** | **7.7%**<br>**(1/13)** | **H5 + H16 (n=1)** |
| Guillemot | 41.3%<br>(19/46) | **2.4%**<br>**(1/42)** | 0/15<br>(0%) | 0%<br>(0/15) | **33.3%**<br>**(6/18)** |  |
| Puffin | 36.3%<br>(16/44) | 0%<br>(0/42) | 0/13<br>(0%) | 0%<br>(0/13) | **7.7%**<br>**(1/13)** |  |

### 3.3 Serology – specific testing for antibodies to different subtypes

Specific testing via haemagglutination inhibition assay was first carried out for the detection of H5 subtype-specific influenza A (Table 2). All kittiwakes, puffins, razorbills and shags had negative H5 HI assay results, while a single guillemot demonstrated a positive titre value (1:16). We subsequently retested samples with sufficient plasma using the newly developed ID Screen® Influenza H5 Antibody Competition 3.0 Multi-species, ID Vet, Montpellier, France. This ELISA assay generated a positive result for H5 antibodies in one puffin, one razorbill, six guillemots and the one shag that had tested positive for generic AIV antibodies. One guillemot demonstrating a positive H5 ELISA was a weak reactor on H5 HI assay (titre 1:2). The rest of these H5 ELISA positive individuals were negative via HI. Despite having the highest prevalence of anti-NP antibody, indicative of previous exposure to AIV H1-H16 (>78% of individuals), no kittiwakes tested positive for H5 antibodies.

Haemagglutination inhibition assay for the detection of H13 and H16 subtype-specific antibodies varied substantially between species (Table 2). Kittiwakes demonstrated high seropositivity to H16 (20/21 of generic AIV positives tested, 95.2%), with four individuals testing concurrently positive for both H16 and H13 antibodies. No antibodies to H13 were detected in any other species. H16 was also found in razorbills and accounted for 4/13 positives detected (30.8%). No shag, puffins or guillemots tested positive for H16. One razorbill tested concurrently for antibodies to H16 and H5.

## 4. Discussion

Empirical data enabling the estimation of key epidemiological parameters is vital in predicting the transmission and persistence of AIV in novel species groups. Quantification of the levels of exposure, the existence of asymptomatic carriers and the presence of potential immunity following exposure and/or recovery is required, accounting for species differences that may co-occur within multi-species assemblages. We found low levels of active, but asymptomatic, AIV infection in individuals (1.6-4.5%), but excluded this as H5N1. We also found clear species variation in antibody prevalence, indicative of previous infection and survival from infection, with seropositivity ranging from 1.1% of shags for antibodies to non-specific AIV antigens to over 70% of kittiwakes, and intermediate levels of detection across auk species (31-41%). Species variation was further detected in relation to the specificity of antibodies to different AIV subtypes. Kittiwakes tested positive only for antibodies to H16 and H13, two AIV subtypes generally associated with infection of gulls^38,56,57^. Despite reports of high mortality among kittiwakes (2022 and 2023), and high levels of AIV NP antibody prevalence observed, we did not detect any H5 antibodies indicative of previous exposure and survival from H5 infection or antigen exposure in this species. In contrast, we found H5 reactive antibodies in guillemots, razorbills, puffins and the single shag that previously tested positive for antibodies to AIV NP. We observed co-occurrence of antibodies to two subtypes in 8.8% of the individuals that were tested for all three subtypes; four kittiwakes had concurrent antibodies to H13 and H16 and a razorbill had concurrent antibodies to H16 and H5. In addition, at least 28 birds (all auks) were antibody positive to an as yet unidentified subtype, suggesting further viral subtypes are likely circulating in the population.

Although we found some evidence of active AIV infection in auk species on the IOM, we have no evidence to confirm that this is H5N1. Previous studies comparing the relationship between the M gene and the HPH5 rRT-PCR assays during UK H5N1 clade 2.3.4.4b HPAIV poultry outbreaks, indicated higher sensitivity in the HPH5 assay. It is highly likely that our M gene positive results relate to H5 LPAIV or another LPAIV. Low levels of active infection are consistent with previously detected levels of LPAIV circulating in migratory wildfowl outside of the current epizootic^58^. Only one individual (razorbill) demonstrated a positive result for high pathogenicity H5 via HPH5 assay. The neuraminidase subtype of this individual remains unknown, with a negative N1 result.

As regards previous exposure to a range of AIV subtypes, our data revealed frequent detection of generic AIV NP antibodies indicative of previous AIV infection with any subtype. The serological data highlighted marked species differences in levels of seropositivity, ranging from extremely high in kittiwakes (>70%) to extremely low in shags (<2%), and at an intermediate level (31-41%) for all three auk species. This mirrors reported patterns in HPH5 mortality, with kittiwakes and shags among the most and least affected species respectively in terms of mortalities attributed to H5N1, both in our own study site and more widely across the UK and Europe^59,60^. Such variation is suggestive of inherent species differences in exposure and susceptibility to AIV within seabird assemblages. Interestingly, the reverse pattern is found in other respiratory viruses such as avian paramyxovirus, which is found at high prevalence in shags but not in other species (*Ravenswater*, *Greco et al, unpublished data*)). Species variation in exposure to infectious virus and susceptibility to infection may arise through inherent species differences in behavioural and ecological differences affecting infection dynamics.

Species differences in social behaviour and in the density, composition and social structure of breeding colonies may generate variation in HPAIV exposure and transmission^61^, as could varying degrees of contact with freshwater or other relevant environments^62^. Similarly, variation in foraging strategy, for example dense feeding aggregations or scavenging behaviour, or in species-specific movement ecology, such as migratory, foraging or prospecting behaviours can introduce further variation in viral dynamics^38,40,63^. Genetic differences in pathogen susceptibility may also occur between-species, particularly in isolated species with historically reduced immune challenge. This is exemplified by the extreme sensitivity to HPAI and limited range of immune gene families observed in the genome of Australian black swans (*Cygnus atratus*) ^64^, particularly when compared to the expansive repertoire of immune genes demonstrated in more resistant species^64,65^. Likewise, differential expression of immune genes, and their corresponding inflammatory responses, can influence susceptibility, resistance and outcome of infection across species^66^. Indeed, it is evident that a complex interaction of factors may drive the dynamics of AIV in each species, or population.

In addition to variation in antibody seropositivity, we found species differences in subtypes to which antibodies were detected. Subsequent to testing for H5, we tested for H16 and H13, as the Laridae family (*Charadriiformes)*, of which the kittiwake is a member, is reported as a natural reservoir for both these subtypes^12,38,67^. We found H16 seropositivity in over 90% of seropositive kittiwakes and H13 in 19%. This observation is consistent with studies prior to the current H5N1 HPAIV outbreak; serological survey of apparently healthy black-legged kittiwakes from Norway, tested in 2009, revealed an antibody prevalence of 71.3% from 80 individuals. HI assay of positive samples from the same study indicated H13 (37.5%) and H16 (81.3%) subtype specificity, with no H5 subtype specific antibodies detected^56^.

Interestingly, although H16 and H13 are almost exclusively found within gull species^38^, we find prevalence of H16 antibodies in razorbill (4/13, 30.8%), a member of the Alcidae family. To the best of our knowledge, serological testing for H13/H16 has not previously been conducted in auks.

Prevalence levels are hence unknown, making cross-population comparisons difficult. Previous studies, including those from Newfoundland and Labrador, Canada^68^; have detected a range of LPAIV in auks, including multiple H1 viruses, and the identification of H3N2 in puffins corroborating the circulation of other subtypes in auk populations^57^. Understanding which subtypes may co-occur is useful in establishing the potential for changes in virus pathogenicity and transmissibility that could occur through reassortment events.

We observed co-occurrence of antibodies to more than one subtype in 8.8% of the individuals that were tested for all three subtypes, highlighting the possible importance of seabirds in the reassortment of AIVs. Viral reassortment between co-infecting AIVs can lead to the emergence of novel viral strains^19^, with substantial genetic diversity noted in circulating H5Nx viruses following recombination and reassortment with LPAIVs^11,39^. Alteration of viral properties via these mechanisms, such as receptor binding specificity or replication rate, can generate viruses with improved evasion of host defences, increased transmissibility or adaptation to new environments and species^69,70^. These adaptations are of particular concern in the reassortment of HPAIV, where the emergence of novel genotypes may have significant consequences for wildlife, domestic animal and human health. For example, during 2022, the reassortment of the AB H5N1 HPAIV genotype with H13 LPAIV, an AIV subtype almost exclusively noted within gull species^38^, produced a novel H5N1 genotype (BB) (see glossary), with widespread circulation and mass mortality in European seabird species^39^. This genotype retained gene segments from gull-specific H13 may explain the increased replication and transmissibility noted within related host species^39^. In addition, while avian-adapted influenza viruses generally replicate and transmit poorly in mammals^19^, the recent spillover of HPAIV H5N1 to over 48 different mammalian species to date^5^ poses a significant risk for the emergence of mammalian-adapted lineages^70^. While evidence for sustained mammalian transmission of clade 2.3.4.4.b H5N1 remains scarce, genetic mutations associated with enhanced mammalian activity are apparent in clade 2.3.4.4.b H5N1 lineages affecting multiple mammalian species within the recent panzootic^5,29,35,36^. If further spillover was to occur to mammalian hosts with co-infecting Influenza A viruses (IAVs), such as the mix of avian, swine and human IAVs, the risk of reassortant strains with zoonotic potential and sustained transmission increases^71,72^.

Despite detecting antibodies at high prevalence levels in kittiwakes, the species with highest mortality from H5N1 HPAIV based on the testing of carcasses in 2022 and 2023, we did not detect active H5N1 viral shedding or H5 antibodies in apparently healthy kittiwakes. There are a number of explanations for this observation. Firstly, if HPAIV was associated with the rapid onset of clinical signs and mortality, proving entirely lethal and leaving no surviving individuals, it is unlikely that we would detect H5 specific antibodies in the apparently healthy individuals sampled. Alternatively, live sampling was completed 12 days before the first confirmed mortality in 2023 and birds may not have yet been exposed to HPAIV. Furthermore, while our study comprises breeding individuals, the majority of dead kittiwakes were found at a single freshwater site, often heavily utilised by non-breeding or sub-adult birds. This could infer higher mortality in these sub-adult or non-breeding components of the population, or that exposure is spatially restricted, creating clusters of infection within the ongoing epizootic. Both are supported by an absence of mortality observed in breeding kittiwakes. We have no knowledge of the antibody status within the sub-adult component of the population, however, inherent differences in the age of individuals could influence viral immunity and the outcome of infection between mature breeding adults and sub-adults within the population, with evidence from mute swans (*Cygnus olor*) indicating the presence of antibodies to an increased number of HA subtypes with age^73^.

Whether the presence of antibodies to any of these subtypes play a protective role remains unclear. Experimental evidence of a relationship between pre-exposure to LPAIV and subsequent H5N1 HPAIV challenge exists across wildfowl, including mallards *(Anas platyrhynchos)* ^74^, wood ducks (Aix sponsa) ^75^ and Canada geese (Branta canadensis) ^41^. These studies reveal reduced mortality and viral shedding in treatment groups inoculated with LPAIV and later challenged with HPAIV when compared to the inoculation of naïve controls with HPAIV. The greatest reduction in clinical signs and viral titres occurs with immunity generated via inoculation with homosubtypic strains of LPAIV (i.e., those of the same/homologous HA subtype, namely H5), yet in some cases a reduction in clinical signs has also been demonstrated with prior exposure to LPAIVs of a different (heterologous) HA subtype^42^. Previous experimental infections in chickens have also demonstrated that cross-subtype protection may occur, where it has been suggested that cell mediated immunity to viral antigens other than the surface HA and NA glycoproteins may have a role. We revealed a high prevalence of H16 antibodies in kittiwakes, yet any potential mechanism for heterologous protection would require further investigation. Furthermore, consideration should also be given to the role of co-occurring and co-infecting pathogens in modulating the outcome of AIV infection, with host differences in pathogen diversity having potential consequences for viral epidemiology via the synergistic or antagonistic effects of co-pathogens^76,77^. Thus, concurrent screening for pathogens beyond AIV is recommended.

Our study also highlights differences in methodological sensitivities, and the need for developing efficient and cost-effective ways of testing for antibodies across a wider range of serotypes, in order to further understand co-occurrence and the potential for reassortment events where different subtypes may be circulating. A higher number of individuals tested positive for H5 specific antibodies when utilising the H5 specific cELISA compared to HI, suggesting that this assay may be more sensitive for detecting H5 antibodies in wild seabirds. While the HI assay is widely used for H5 subtyping of antibodies, including in studies of other overlapping seabird populations within the same time frame^8^, the sensitivity of HI can be impacted by the similarity of the genotype of the virus from which the test antigen is derived^78^. Within our study we utilised test antigen H5N1 HP (A/Chicken/Wales/053969/21), considered homologous to the current HPAIV outbreak, although some minor antigenic variation has been observed in the clade 2.3.4.4b subtype since the outbreak started^79^.

Viral circulation is restricted by a relatively short period of active AIV infection, whereby viral shedding normally resolves (in non-fatal AIV infections) by around 10 days post infection^80,81^, as detected by rRT-PCR. Serological detection of antibodies is less time critical, with experimental evidence from black-headed gulls (*Chroicocephalus ridibundus*) indicating the persistence of AIV H16 subtype specific antibodies over eleven months^81^, emphasising the advantages of combined surveillance via both molecular and serological methods. Further, in individuals testing positive for AIV RNA, three were detected on oropharyngeal swabs, and two on cloacal swabs. ^82,83^. We detect no individual concurrently positive on oropharyngeal and cloacal samples, supporting the need for the combined swabbing approach recommended by previous studies^67,83^.

In summary, data informing the epidemiology of AIV in co-occurring species groups is vital in predicting the ongoing viral transmission, persistence and consequences of H5N1 clade 2.3.4.4b HPAIV, and the range of AIVs, that circulate within wild avian populations. There is an urgent need to consider the role of seabirds in the ongoing epidemiology of HPAI due to both their long-distance migratory behaviour, which has the capacity to transmit and increase exposure to a diverse range of viral variants across a broad geographic range, and due to the wide range of habitats used. This includes both coastal and urban environments enhancing the likelihood of transmission at the interface of wild and domestic species. We therefore promote the need for wider studies beyond disease surveillance, including the identification of co-circulating viral subtypes and antibodies within multi-species assemblages impacted by H5Nx HPAIV. This is imperative in providing data for the multi-disciplinary understanding and prediction of risks associated with viral recombination and emergence within closely associated populations, and within the wider context of One Health.

## Acknowledgments

We would like to thank Ruth E. Dunn and Eve Merrall for their help in collecting this data. We thank NatureScot for granting permission to work on the Isle of May, and their staff and volunteers for help in the field. FG was funded by a Natural Environment Research Council (NERC) Doctoral Training Partnership grant (NE/S007407/1), HR by NERC Doctoral Training Partnership grant NE/L002558/1, EC by NERC grants NE/V001779/1: A direct test of the impact of infection on animal migration, NE/X018261/1: Creating an urgently needed infrastructure for avian influenza monitoring in wild birds and NE/Y001591/1 ECOFLU: Understanding the ecology of Highly Pathogenic Avian Influenza in wild bird populations, FRR and CD by NERC grant NE/V001779/1: A direct test of the impact of infection on animal migration. UKCEH authors were supported by Natural Environment Research Council Award number NE/R016429/1 as part of the UK-SCAPE Programme Delivering National Capability. ST, MF, SMR, MJS, JJ and ACB were funded by the UK Department for the Environment, Food and Rural Affairs (Defra) and the devolved Scottish and Welsh governments under grants SE2213, SE2227, SV3400, SV3006, and SE2230. MJS, JJ and ACB were also part funded by the Biotechnology and Biological Sciences Research Council (BBSRC) and Department for Environment, Food and Rural Affairs (Defra, UK) research initiative ‘FluMAP’ [grant number BB/X006204/1] and ‘FluTrailMap’ [grant number BB/Y007271/1].

## Author contributions

F.G., E.C., H.R. and S.B. conceived the study.

F.G., H.R., C.D. F.R-R., S.B., E.C., M.N., J.H., E.T., E.B., F.D., D.S. collected the data

J.P., E.P., D.S. collated mortality data

F.G., H.R., S.T., M.S. conducted the analyses F.G., E.C., S.B. wrote the paper

H.R., F.R-R., C.D., M.N., J.H., E.T., E.B., F.D, G.G., D.S., J.P., E.P., S.T., M.S., A.C.B., J.J., S.R., M.F. contributed to discussion and comments on the paper

## Data availability statement

Datasets generated during this study are available from the corresponding author on reasonable request.

## Additional Information Competing Interests

The authors declare no competing interests.

