## Supplementary material for "Asymptomatic infection and antibody prevalence to co-occurring avian influenza viruses vary substantially between sympatric seabird species following H5N1 outbreaks"

**†** Shared last authorship

*

**Supplementary information**

**Mortality events – summer 2023**

Two kittiwake carcasses swabbed on 26^th^ June 2023 tested positive for AI viral RNA. A total of 871 kittiwake deaths were subsequently reported as suspected H5N1, including 190 fledglings and 681 sub-adults/adults. These deaths were predominantly concentrated in a single geographical location around a freshwater loch. Mortality of unknown cause was also recorded in 30 individuals of other species, namely herring gull (n=6), lesser-black backed gull (n=3), puffin (n=17) and eider (n=4) (*NatureScot, unpublished data*). These latter records of mortality across the season do not appear atypical in level in for these species.

***Supplementary Table S1:*** *Predominant species assemblage on the Isle of May NNR and relevant UK Bird of Conservation Concern status. Where available, breeding population estimates for each species on the IOM are presented for the year prior to sampling (2022): AON = apparently occupied nests, AOS = apparently occupied sites, AOB = apparently occupied burrows. Where data is not available for 2022, the previous census data is given – kittiwakes (2021) and puffins (2017). With the exception of eiders where the figure for breeding males is unknown, these values represent breeding pairs and the breeding population of individuals will be double these figures. Note that non-breeders present are not included in these estimates. Also noted is the UK breeding population as a percentage of the international population. Finally, the percentage change in breeding population size of each species in the designated Forth Islands Special Protection Area between the seabird population census’ Seabird 2000 (1998-2002) and Seabirds Count (2015-2021) is given. These population changes pre-date the HPAI H5N1 clade 2.3.4.4b epizootic*, *with some species already showing clear declines prior to the emergence of HPAIV.*

| **Species** | **Isle of May Species Estimates 2022^1,2^** | **UK Birds of Conservation Concern 5^3,4^** | **UK population as % of international population^5^** | **Forth Island SPA % breeding population change^6^** |
| --- | --- | --- | --- | --- |
| Puffin  (*Fratercula arctica*) | 39,200 AOB  (*2017 census*) | Red | 9.6% | -40% |
| Black-legged kittiwake (*Rissa tridactyla*) | 5193 AON  (*2021 census*) | Red | 8% | -22% |
| Herring gull  (*Larus argentatus*) | 5168 AON | Red | 12.1% | -14% |
| Arctic tern  (*Sterna paradisaea*) | 578 AON | Red | 3.1% | -8% |
| Great black-backed gull (*Larus marinus*) | 116 AON | Red | 9.6% | NA |
| European shag  (*Gulosus aristotelis*) | 467 AON | Amber | 34.1% | -30% |
| Lesser black-backed gull (*Larus fuscus*) | 1739 AON | Amber | 38.4% | -31% |
| Common guillemot (*Uria aalge*) | 17,318 AOS | Amber | 12.9% | -27% |
| Northern fulmar (*Fulmarus glacialis*) | 321 AON | Amber | 8% | NA |
| Razorbill  (*Alca torda*) | 4381 AOS | Amber | 20.2% | +23% |
| Common eider (Somateria mollissima) | 715 nesting females | Amber | NA | NA |

***Supplementary Table S2*:** *Number of total positive (Cq ≤36.0) or borderline (Cq >36.0, <40.0) swab samples per number of individuals for each species. M-gene assay refers to the detection of nucleic acid from all sixteen generic HA AIV subtypes, while H5HP assay is specific for nucleic acid of HPAI H5Nx pathotype. Species prevalence for positive and borderline results is given in brackets, unless zero detection*. *Both positive and borderline results are displayed in* ***bold****. Range of infected individuals gives a minimum and maximum percentage of positive viral detection of AIV (all subtypes) per samples for each species. The minimum value indicates confirmed positive individuals only, while maximum value combines positive and borderline individuals. Note that, one individual puffin tested borderline across two assays, so the total number of borderlines plus positive results in puffins is n=6. Our estimate of positive individuals across the entire set sampled ranged from 1.6-4.5% (5/309 to 14/309)*.

| **Species** | **M-gene** | | **H5 HP** | | **N1** | | **Range of infected individuals** |
| --- | --- | --- | --- | --- | --- | --- | --- |
|  | **Positive** | **Borderline** | **Positive** | **Borderline** | **Positive** | **Borderline** |  |
| Shag | 0/98 | 0/98 | 0/98 | 0/98 | NA | NA | NA |
| Kittiwake | 0/74 | 0/74 | 0/74 | 0/74 | NA | NA | NA |
| Puffin | **1/45 (2.2%)** | **3/45 (6.7%)** | 0/45 | **3/45 (6.7%)** | 0/3 | **1/3** | 2.3% (1/45) –  13.3% (6/45) |
| Guillemot | **3/44**  **(6.8%)** | **4/44**  **(9.1%)** | 0/44 | 0/44 | **1/7** | **2/7** | 6.8% (3/44) – 15.9% (7/44) |
| Razorbill | 0/48 | 0/48 | **1/48 (2.1%)** | 0/48 | 0/1 | 0/1 | 2.1% |

***Supplementary Table S3:*** *Results breakdown across assay type in individuals reported positive (Cq ≤36.0) or borderline (given here, Cq 36.01 – 36.9) for viral nucleic acid via rRT-PCR (positives in* ***bold****). Cq value or negative result given, NA = assay not performed. Swab type CL= detection on cloacal swab, CH= detection on choanal swab. An indication of the final subtyping achieved for each positive sample is given: HxNx = AIV of unknown subtype; H5Nx = high pathogenicity H5, unconfirmed neuraminidase subtype; HxN1 = confirmed N1, unconfirmed haemagglutinin subtype. Associated competitive ELISA assay (H1-H16 and H5 specific) and haemagglutination inhibition (HI) results also given for each individual.*

| **Species** | **Sample date** | **Swab type** | **Assay type** | | | **Indicated subtype of positive samples** | **ELISA**  **(H1-H16)** | **ELISA**  **(H5)** | **H5 HI** |
| --- | --- | --- | --- | --- | --- | --- | --- | --- | --- |
|  |  |  | **M-gene** | **H5 HP** | **N1** |  |  |  |  |
| **Puffin** | **06/07/2023** | OP | **35.64** | No Cq | No Cq | HxNx | Negative | Not tested | Negative |
|  |  | C | No Cq | No Cq | NA |  |  |  |  |
| **Guillemot** | **21/06/2023** | OP | No Cq | No Cq | NA | HxNx | **Positive** | **Positive** | Negative |
|  |  | C | **34.38** | No Cq | 36.43 |  |  |  |  |
| **Guillemot** | **21/06/2023** | OP | **35.85** | No Cq | No Cq | HxNx | Negative | Not tested | Negative |
|  |  | C | No Cq | No Cq | NA |  |  |  |  |
| **Guillemot** | **22/06/2023** | OP | No Cq | No Cq | NA | HxN1 | **Positive** | **Positive** | Negative/ |
|  |  | C | **34.28** | No Cq | **35.02** |  |  |  | Weak reactor |
| **Razorbill** | **17/06/2023** | OP | No Cq | **34.97** | No Cq | HP H5Nx | Negative | Not tested | Negative |
|  |  | C | No Cq | No Cq | NA |  |  |  |  |
| Puffin | 11/06/2023 | OP | No Cq | No Cq | NA |  | Not tested | Not tested | Negative |
|  |  | C | 36.36 | 36.55 | 38.25 |  |  |  |  |
| Guillemot | 21/06/2023 | OP | 36.62 | No Cq | No Cq |  | Negative | Not tested | Negative |
|  |  | C | No Cq | No Cq | NA |  |  |  |  |
| Guillemot | 21/06/2023 | OP | 36.4 | No Cq | No Cq |  | Positive | Negative | Negative |
|  |  | C | No Cq | No Cq | NA |  |  |  |  |
| Guillemot | 21/06/2023 | OP | No Cq | No Cq | NA |  | Negative | Negative | Negative |
|  |  | C | 36.89 | No Cq | 36.43 |  |  |  |  |
| Guillemot | 21/06/2023 | OP | 36.75 | No Cq | No Cq |  | Positive | Negative | Negative |
|  |  | C | No Cq | No Cq | NA |  |  |  |  |
| Puffin | 06/07/2023 | OP | 36.91 | No Cq | NA |  | Positive | Negative | Negative |
|  |  | C | No Cq | No Cq | NA |  |  |  |  |
| Puffin | 29/06/2023 | OP | No Cq | 37.5 | NA |  | Negative | Not tested | Negative |
|  |  | C | No Cq | 39.09 | NA |  |  |  |  |
| Puffin | 29/06/2023 | OP | No Cq | No Cq | NA |  | Negative | Not tested | Negative |
|  |  | C | 38.32 | No Cq | NA |  |  |  |  |
| Puffin | 06/07/2023 | OP | No Cq | No Cq | NA |  | Negative | Not tested | Negative |
|  |  | C | No Cq | 38.68 | NA |  |  |  |  |
